## Supplementary Figures for "Representations in human primary visual cortex drift over time"

Laboratory of Brain and Cognition
National Institute of Mental Health, NIH,
Bethesda, MD, USA

This file includes Figures S1 to S5

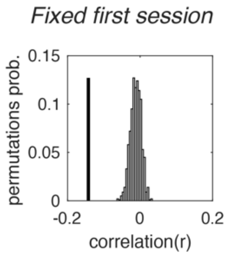


***Fig S1.*** *Null distribution of correlation between $R^2$ and $\Delta$session, while keeping the first session fixed in all permutations. Drift is still significant ($p<0.001$), indicating that it is not driven solely by different responses in the first session.*

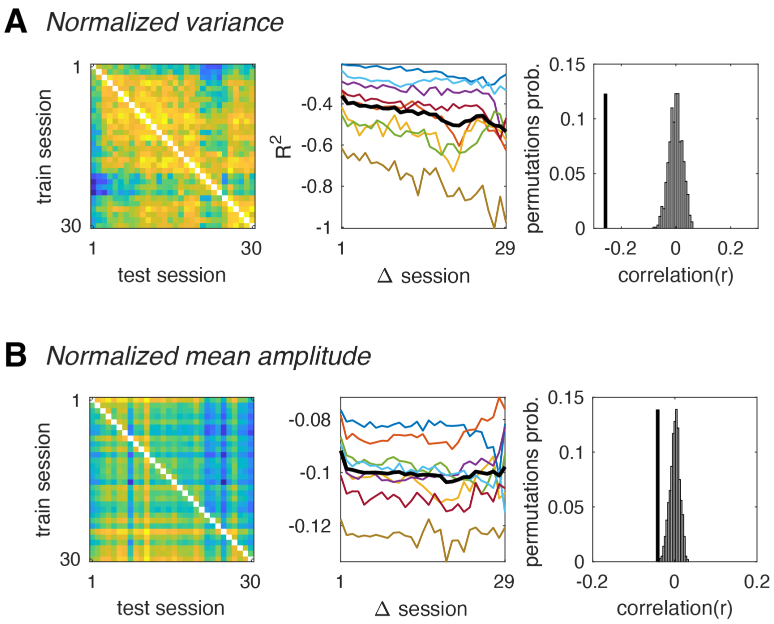


***Fig S2. A**. Cross-session generalization after normalizing each session's variance. Left, goodness-of-fit matrix after normalizing each voxel's response variance within each session. Center, Mean $R^2$ as function of number of intervening sessions between train and test sessions. Model predictive power decreases with time, indicating representational drift (r=-0.26, p<0.001). gray lines, individual subjects; thick black line, mean across subjects. Right, Black vertical line, empirical correlation between goodness-of-fit and number of intervening sessions. Gray histogram, null distribution of correlation values. **B**. Cross-session generalization after subtracting each session's mean response amplitude (i.e. mean beta). Left, goodness-of-fit matrix after subtracting each voxel's mean response within each session. Center, Mean $R^2$ as function of number of intervening sessions between train and test sessions. Predictive power of V1 models still decreases significantly with time (r=-0.04, p=0.001), but much less than without subtracting the mean.*

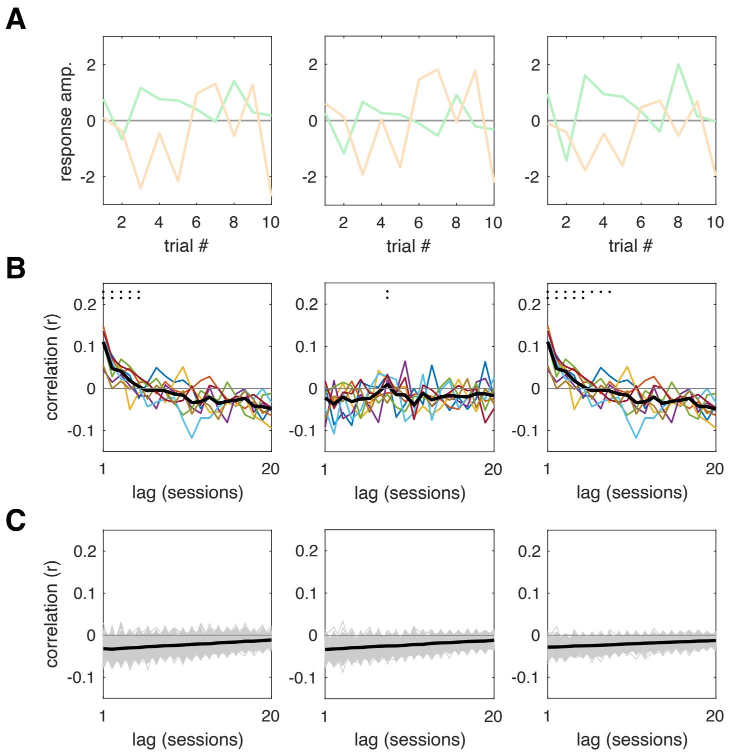


***Fig S3.*** *Mean response amplitude gradually changes across sessions.* ***A****. Schematic illustration of original responses (left), responses after removing mean (center), and responses after normalizing variability (right).* ***B.*** *Autocorrelation of voxelwise mean beta, black dots indicate values significantly above zero (1 dot, p<0.05; 2 dots, p<0.01; uncorrected for multiple comparisons). Autocorrelation values are significantly above zero, up to a lag of 7 sessions, but only for original responses (left), and after normalized STD (right), but not after removing the mean response amplitude (center).* ***C.*** *Null distribution of autocorrelation values. Black line, mean across 1000 permutations; gray lines, values for individual permutations.*

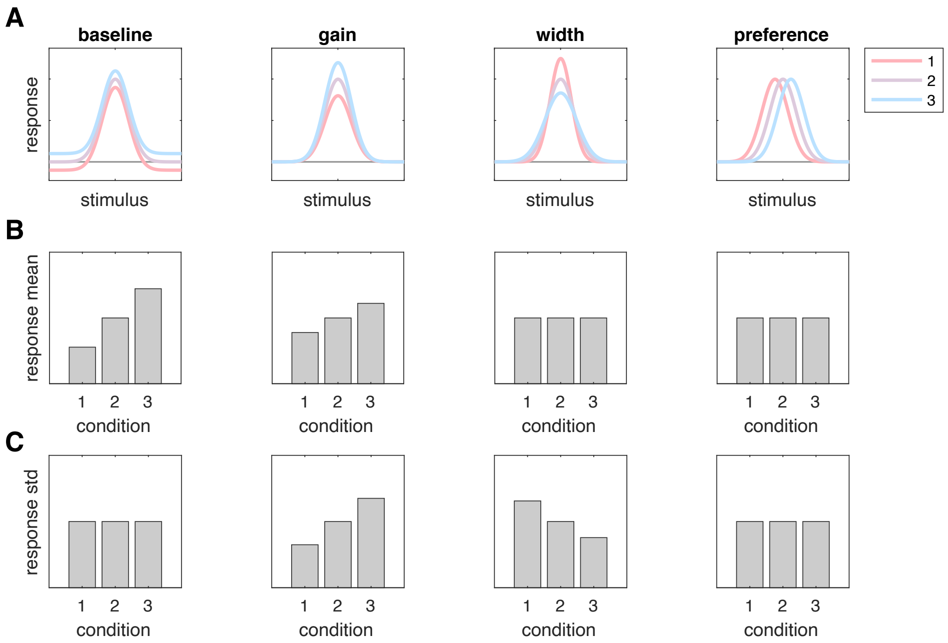


***Fig S4.*** *Hypothetical changes in voxel tuning underlying drift.* ***A.*** *From left to right, possible changes are changes in: baseline, gain, tuning width, and stimulus preference. Three colored tuning curves are illustrated with different values for each possible tuning change.* ***B.*** *Mean response amplitude for each of the possible tuning curves. Changes in baseline and gain affect the mean, while changes in tuning width and preference do not.* ***C.*** *STD of response amplitudes for each of the possible tuning curves. Changes in gain and tuning width affect response STD, while changes in baseline and preference do not. These simulations suggest that only an additive baseline shift would affect the mean response amplitude, without a concomitant change in the variance of responses, consistent with the drift that we observed. Note that normalizing the mean amplitude cannot compensate completely for changes in the baseline, since each session included a different set of stimuli, and therefore was likely to evoke a different mean amplitude even if the baseline remained the same.*

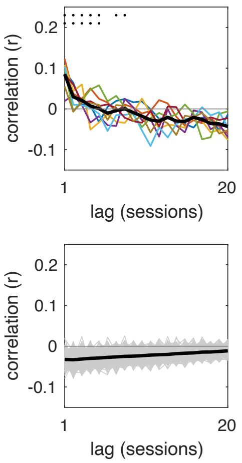


***Fig S5.*** *The weight of the model's constant component changes gradually across sessions. Top row: autocorrelation of the constant component. Black dots indicate values significantly above zero (1 dot, p<0.05; 2 dots, p<0.01; uncorrected for multiple comparisons). Bottom row: Null distribution of constant weight autocorrelation values. These results confirm that, similar to response amplitudes of individual voxels, baseline response values as captured by the image-computable model gradually accumulate changes across sessions and are not simply fluctuating around a fixed mean.*
